# Sex differences in exploration–exploitation strategies during home-cage decision making

**DOI:** 10.64898/2026.04.02.716124

**Authors:** Chantelle L. Murrell, Alex A. Legaria, Katie B. McCullough, Andrew Nwacha, Monsurat O. Nasiru, Sebastian Alves Ferreira Dias, Rebecca Chase, Mason R. Barrett, Matt Gaidica, Naoki Hiratani, Meaghan C. Creed, Joseph D. Dougherty, Susan E. Maloney, Alexxai V. Kravitz

## Abstract

The exploration-exploitation trade-off refers to the conflict between using known strategies that reliably yield reward (exploitation) and sampling uncertain options that might yield better outcomes (exploration). Dysregulation of this balance is implicated in neuropsychiatric disease, and while sex differences in this balance have been described, the biological bases remain unclear. To quantify sex differences in this trade-off, we tested mice (n=74 male, 62 female) on four home-cage based foraging tasks with an operant pellet dispensing device, Feeding Experimentation Device 3 (FED3). Mice completed the tasks continuously over multiple days and the tasks were their only source of food. Across multiple tasks, males showed higher win-stay behaviour than females, indicating greater exploitation of previously rewarded actions, an effect that was modest in size but highly significant. Power analyses revealed that >30 mice per sex were needed to detect these modest but significant sex differences with 80% power. No consistent sex differences were observed in pellet intake, suggesting that differences in exploitation did not reflect differences in hunger drive or demand for pellets. Exploitation is a more efficient strategy when environmental parameters are fixed, while exploration can be more advantageous when parameters such as reward locations are changing and uncertain. We tested this idea by re-running our mice in a probabilistic foraging task, where actions led to uncertain probabilities of reward. While males continued to show higher levels of win-stay behaviour on this task, this no longer led to increases in accuracy. Behavioural modelling also supported this framework, demonstrating that stronger win-stay behaviour was most advantageous in deterministic models, and less advantageous in probabilistic models. Together, our findings demonstrate that male and female mice have small but significant differences in their exploration-exploitation balance, which leads to more accurate foraging in certain, but not uncertain, environments.

## Intro

Animals must make decisions under uncertainty to survive in changing environments. For example, when animals leave the safety of their nest, they need to decide whether to forage for food in locations where they previously obtained food, or to explore new locations that might contain better outcomes. This has been framed as a balance between “exploitation” versus “exploration” (Speekenbrink and Konstantinidis 2015; Sutton and Barto 1998). Should animals exploit known information to reap certain rewards? Or should they devote time and energy to exploration to learn new things about the environment? Difficulties in balancing these processes have been implicated in neurodevelopmental and neuropsychiatric disorders. For example, too much exploration may be linked to impulsive behaviour and indecision (Dubois and Hauser 2022), while an extreme level of exploitation may be emblematic of problematic compulsions in obsessive compulsive disorders, substance use disorders (Jami et al. 2025), or restricted interests/repetitive behaviours in autism (Goetmaeckers et al. 2025; Solomon et al. 2011).

Sex differences in the balance of exploitation versus exploration have been reported in both humans and rodents, although the literature on which sex has a stronger drive for exploitation is inconclusive and potentially contradictory (Chen et al. 2021; van den Bos et al. 2012; Kalhan et al. 2026). These observations suggest that exploration–exploitation strategies may serve as a behavioural phenotype through which underlying sex-specific vulnerabilities to neuropsychiatric disorders are expressed for example, males show higher rates of diagnosis for autism (Loomes et al. 2017; Werling and Geschwind 2013) and substance use disorders (Becker and Hu 2008; Van Etten et al. 1999; McHugh et al. 2018), while adult females have higher rates of OCD (Mathis et al. 2011; Mathes et al. 2019).

To address this, we tested whether male or female mice had a stronger drive for exploitation, and we quantified how this impacts the efficiency of their foraging behaviour in multiple food-rewarded operant tasks. The primary task we used for this was the two-armed bandit task, delivered via the FED3 device (Fisher et al. 2024). Two-armed bandit tasks require mice to choose between two options that deliver rewards with different probabilities. Previous literature reports used this task on the FED3 device to examine reversal learning and flexible cognition, however none have evaluated the strategies used between male and female mice (Yao et al. 2026; Murdaugh et al. 2024; Li et al. 2026; Conn et al. 2024). Examining how faithfully mice direct their behaviour at the higher reward option can reveal how well they understand the task. Examining how current choices are influenced by prior choices can further reveal the strategies mice are using to solve the task. For instance, do they choose the option that led to a reward on the previous trial (ie: exploitation)? Or do they still test out the non-rewarded option (ie: exploration)? Finally, reversing the reward probabilities between the two options can reveal how flexible and quickly mice can update their behaviour when the environment changes.

Many prior studies examining exploration and exploitation were conducted in male animals, precluding comparisons between the sexes. Only in recent years have researchers challenged the assumption of reduced variability in males (Beery 2018; Kaluve et al. 2022), and emphasized the importance of sex as a biological variable (Clayton 2016; Lee 2018; Grissom et al. 2024). Due in part to the limited number of studies that directly compare male and female animals, it remains unclear whether males or females have a higher drive for exploitation or exploration. It has also become clear that the specific task structure can also influence these processes, making it challenging to compare between different tasks.

For example, in a “restless” variant of the two armed bandit task, Chen *et al*., reported that although male and female mice achieved similar accuracy on the task, males displayed a greater tendency to explore whereas females were more likely to exploit known information (Chen et al. 2021). In contrast, other studies have reported evidence for greater exploitation in males: male rats showed an increased win-stay strategy (i.e., exploitative, or *stick with what works*) compared to females in both a rodent Iowa Gambling task (van den Bos et al. 2012), and in a probabilistic reversal learning task (Kalhan et al. 2026). Still other studies have not detected sex differences in exploitation in reversal learning paradigms (Harris et al. 2021; Chowdhury et al. 2019; Murdaugh et al. 2024). The lack of agreement of prior literature may reflect differences in task structure between studies, or small group sizes that are not adequately powered to reliably detect differences in decision-making strategies between males and females (Button et al. 2013).

Here, we sought to quantify sex differences in exploitation between male and female mice. To obtain a sufficiently large sample size to resolve this question, we relied on high-throughput behavioural assays completed in home cages using the Feeding Experimentation Device 3 (FED3) (Matikainen-Ankney et al. 2021). In this way, mice could earn all their daily food from the tasks while remaining in their home-cages, increasing throughput and reducing the impact of experimenter interventions and other disruptions on behaviour (Huzard et al. 2026). We tested mice on four tasks designed to assess learning, food demand, and motivation. Across these four tests, we detected small but significant differences in exploitation drive, such that male mice were more likely to exploit known information, which made them more efficient foragers in tasks with deterministic outcomes, but not in tasks with less certain outcomes.

## Results

One hundred and thirty-six C57BL/6J mice (62F/74M) between 8-16 weeks of age were tested on four home-cage tasks, taking place across 2 weeks (Figure 1). The tasks were programmed into a FED3 (Matikainen-Ankney et al. 2021), which was placed in the cage with the mice to collect continuous data.

**Figure 1.**
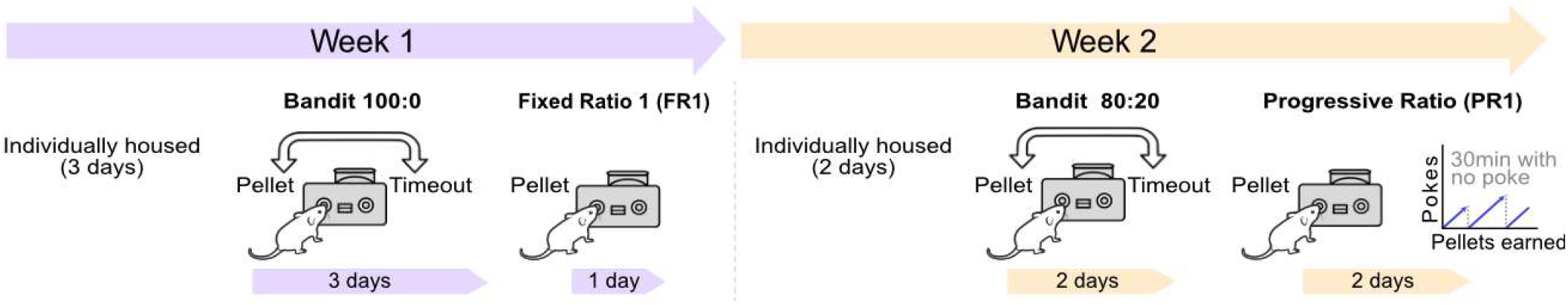
Mouse behavioral training took place over two weeks. In Week 1, naïve mice were individually housed for 3 days and proceeded to complete the Bandit 100:0 task (3 days), and the FR1 task (1 day). In Week 2, mice started with no task over the weekend and then completed the Bandit 80:20 task (2 days), and the PR1 task (2 days).

Mice were single housed and earned all their food from these tasks in a “closed economy” design (Collier 1989).

The first task was a two-armed “Bandit 100:0” task, where one nose-poke resulted in a pellet while the other resulted in a 10s timeout (Figure 2A). Every 20 pellets, these contingencies switched so that the previously inactive nose-poke delivered a pellet, while the previously active nose-poke resulted in a timeout. The bandit task was run for 3 days straight, with females earning an average of 176 ± 3 pellets per day, while males earned an average of 179 ± 3 pellets per day (unpaired t-test, *p* = 0.76). As such, mice completed ∼8-9 contingency switches per day across the 3 days. (Figure 2B). On average, we did not observe a systematic a side bias in this task, and across all mice their individual bias was within ±10% of 50%. In addition, individual levels of poke bias did not significantly correlate with accuracy in the Bandit 100 and FR1 tasks. Male mice were slightly but significantly more accurate than female mice on this task, meaning they more often poked on the side that was paired with the pellet (M = 63.9 ± 0.6%, F = 60.6 ± 0.5%, unpaired t-test *p* = 0.001, Figure 2C). We therefore compared the decision-making strategies of male and female mice on this task, to test if differences in exploitation might relate to these differences in accuracy. We parsed trials into a “win-stay” and “lose-stay” decisions. “Win-stay” meant they returned to the same nose-poke following a rewarded trial, which reflects exploitation. “Lose-stay” is the converse of “lose-shift” and describes trials where mice returned to the same nose-poke following a time-out (see Methods). Males had a significantly higher percentage of win-stay decisions (M = 82 ± 0.7%, F = 77.1 ± 0.9%, Figure 2D, unpaired t-test *p* = 0.001), with no difference in lose-stay decisions (M = 53.1 ± 0.8%, F = 52.5 ± 0.9%, Figure 2E, unpaired t-test *p* = 1.0). We considered if the higher accuracy of males might reflect a greater rate of learning the task. To assess this, we calculated accuracy separately for each day of the task. Mixed-model ANOVA showed effect of Sex (F[1,134]=16.5, p=0.0001), Day (F[2,268]=157, p<0.0001), but no Interaction (F[2,268]=0.49, p=0.61). This indicates that the sex differences we observed are a stable feature of the mice, and do not reflect a higher learning rate of male mice on this task. Finally, we fit a logistic regression model to predict the current choice using their choices on the previous five trials. In both males and females, a rewarded choice on the previous trial was a strong predictor of their choice next trial (Odds ratios for M: 2.89 ± 0.14, F: 2.24 ± 0.13, Figure 2F), although this prediction was stronger for males. In both sexes, further trials back had a minimal impact on the current choice (Figure 2F). We conclude that both male and female mice exhibit strong win-stay behaviour in this task, but males have higher levels of win-stay and accuracy than females.

**Figure 2.**
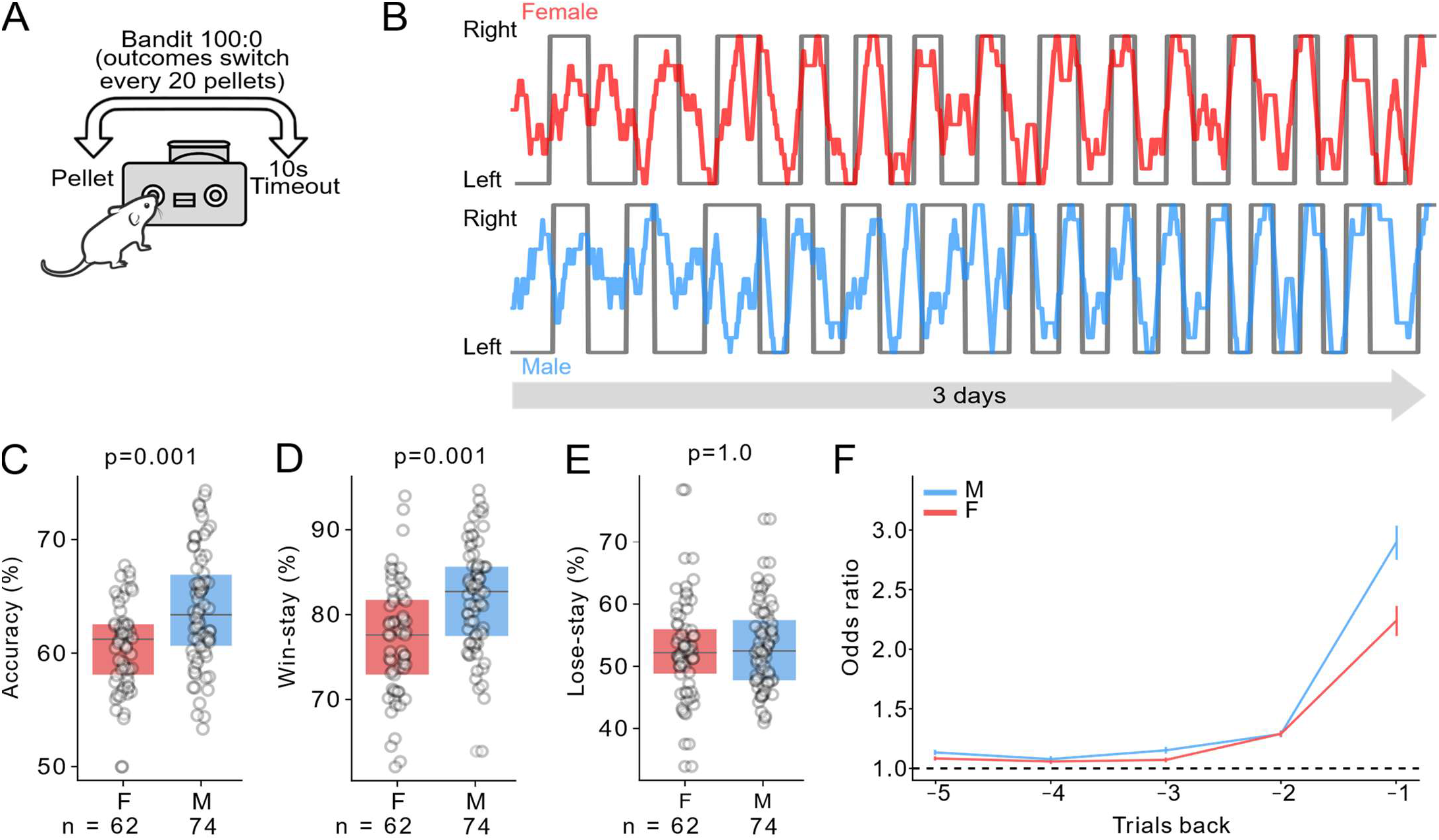
A) Schematic of the Bandit 100:0 task. B) Example of mouse behaviour, each datapoint represents the average choice proportion of 10 trials in a rolling window (F red M blue) with the grey trace indicating the true reward probability in the Bandit 100:0 task. C, D, E) Accuracy, win-stay and lose-stay probability of females vs males (n = 62F / 74M). F) Logistic regression predicting the current choice from the previous 5 choices. P-values were the result of unpaired 2-tailed t-tests, Bonferroni corrected.

Our home-cage based training caused mice to complete trials around-the-clock, meaning we could test if accuracy or decision making strategies differed across the circadian cycle. As expected, both male and female mice took more pellets in the night than in the day (M = 51 ± 2 (day) vs 129 ± 2 (night), F = 46 ± 1 day vs 129 ± 3 night, Figure 3C, paired t-tests *p* < 0.001). However, there were no significant differences in accuracy (M = 63.4% ± 0.8% (day) vs 64.3% ± 0.6% (night), F = 60.2% ± 0.8% (day) vs 60.9% ± 0.5% (night), Figure 3D, paired t-test *p* = 1.0) or win-stay behaviour (M = 82.3% ± 0.8% (day) 81.9% ± 0.8% (night), F = 77.9% ± 1.1% (day) vs 77% ± 0.9% (night), Figure 3E, paired t-test *p* = 1.0) between the day and night, for male or female mice. This suggests that the win-stay strategy is not influenced by the time of day, nor the rate of pellet intake.

**Figure 3.**
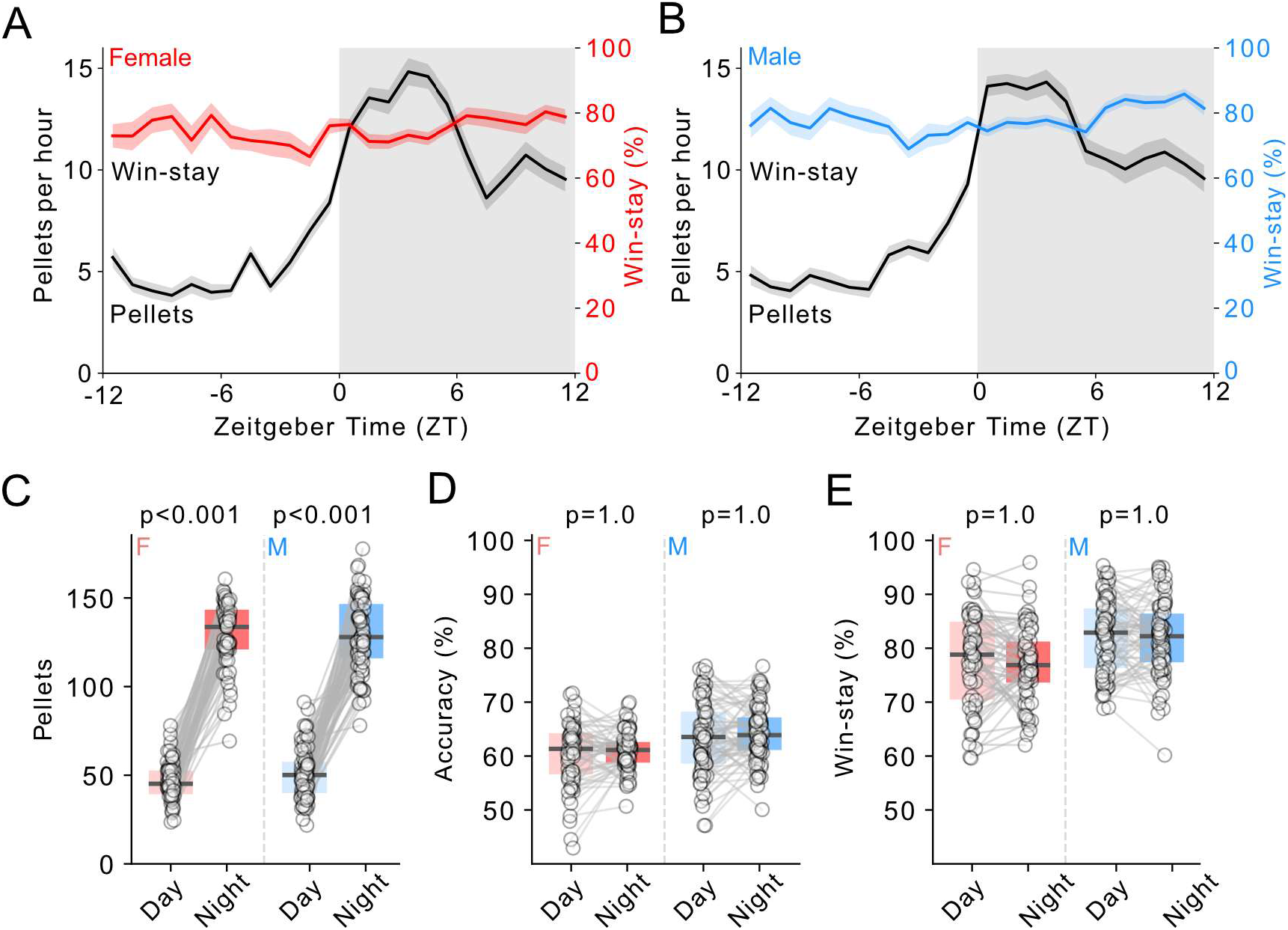
A,B) Win-stay per hour (red/blue) and pellets per hour (black) in female (A) and male (B) mice. C, D, E) Pellets, accuracy and win-stay in the day (inactive phase) and night (active phase) for females (red) and males (black). P-values were the result of paired 2-tailed t-tests, Bonferroni corrected.

We next tested if the difference in win-stay strategy in male mice might reflect a general sex difference observable even outside the Bandit task. Directly following the third day of the Bandit task, mice were trained for an additional day on a Fixed-Ratio 1 (FR1) task, where the Left port remained active for one day without switching and the right port delivered no consequence (Figure 4A). Mice rapidly learned to stop switching and direct most of their poking to the active side (Figure 4B). Male mice took more pellets on this task compared to female mice (M = 187 ± 3, F = 166 ± 3, unpaired t-test *p* < 0.001), and achieved a higher accuracy (% of pokes on the active side) (M = 74.5 ± 1%, F = 66.7 ± 1%, Figure 4B, C, unpaired t-test *p* < 0.001). Male mice also had a higher percentage of win-stay trials (M = 92.4 ± 0.5%, F = 86.9 ± 1%, Figure 4D, unpaired t-test *p* < 0.001). This suggests that the increased propensity of male mice to repeat previously rewarded actions was not unique to the Bandit task but reflected a more general trait of male mice.

**Figure 4.**
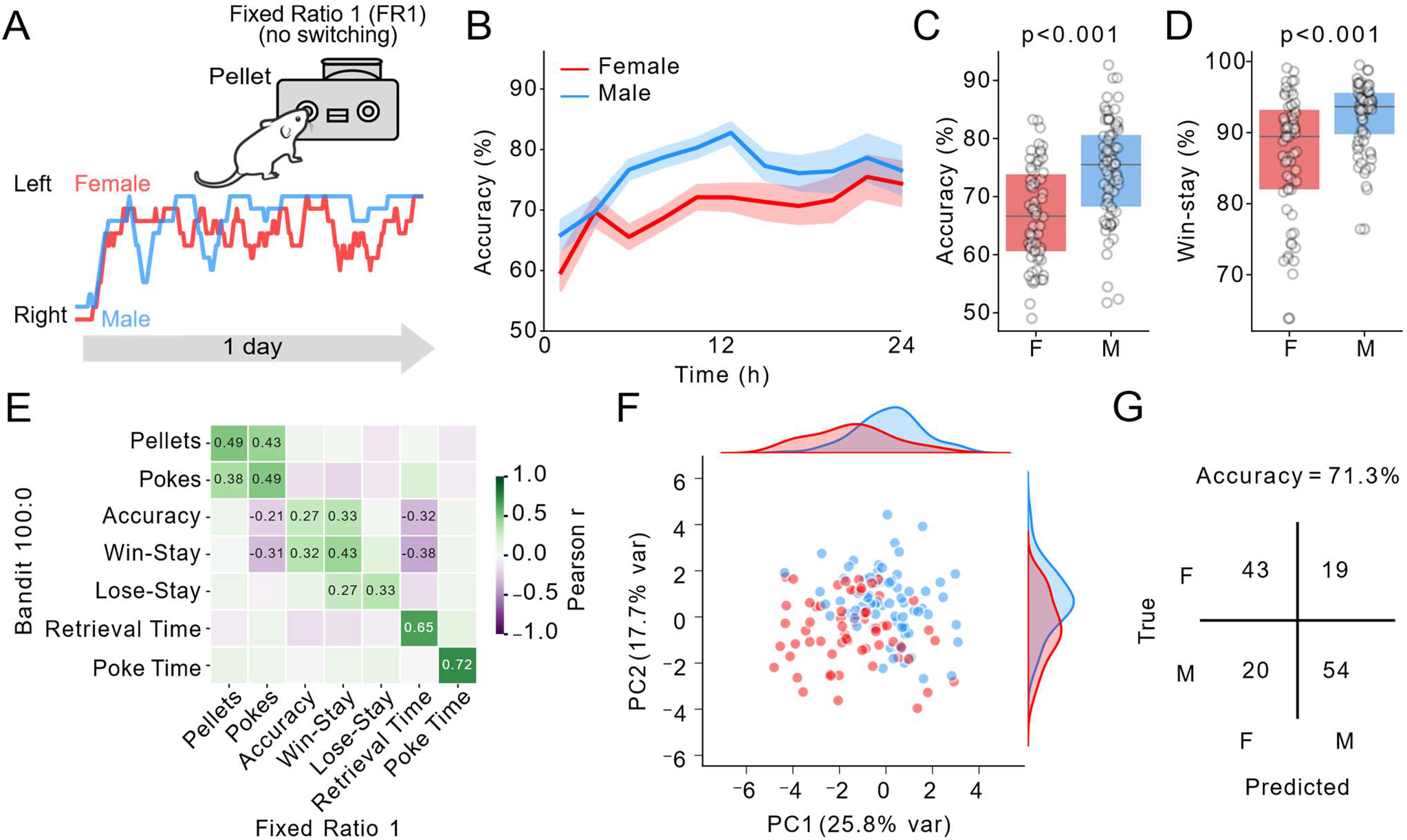
A) Schematic of the fixed ratio 1 (FR1) task with example traces of a single female and male trial (F red, M blue). B, Accuracy over 24 hours, binned in 1hr bins. C, D) Accuracy and win-stay probability of females vs males (n = 62F / 74M). E) Significant correlations of the same metrics between the Bandit 100:0 and FR1 task. F) PCA analysis of 14 behavioural features from Bandit 100:0 and FR1 tasks cluster male (blue) and female (red) mice. G) Confusion matrix predicting sex from our behavioural data using accuracy, win-stay, total pokes and retrieval time from Bandit 100:0 and FR1 tasks. P-values were the result of unpaired 2-tailed t-tests, Bonferroni corrected.

To compare the stability of individual mouse behaviour between Bandit and FR1, we performed correlational analyses that revealed positive correlations for pellets earned, number of pokes, accuracy, win-stay, lose-stay, retrieval time, and poke-time between the tasks (Figure 4E). We then performed principal component analysis (PCA) on these features, revealing that PC1 provided relatively strong separation between male and female mice (Figure 4F). Among the features contributing most strongly to PC1 across the two tasks were accuracy, win-stay, pokes and retrieval time (which was faster for males). We therefore trained a logistic regression model to predict sex from these four features from the Bandit 100:0 and FR1 task, summarizing its performance at predicting sex with a confusion matrix (Figure 4G). The regression model classified mice with an overall accuracy of 71.3%, demonstrating that this set of behavioural features was sufficient to distinguish male from female mice with moderate accuracy Prior literature has shown that win-stay behaviour is most beneficial in deterministic environments, where reward reliably identifies the correct response, but is less useful in probabilistic environments in which unrewarded trials can occur despite correct choices and reward contingencies may reverse over time (Swanson et al. 2022). To explore the possible impact of win-stay on accuracy in different environments, we modelled a win-stay lose-shift (WSLS) agent whose choices were determined by win-stay, lose-shift, and a ‘lapse’ parameter (Le et al. 2023). The lapse parameter describes disengagement or inattention and was therefore programmed to choose randomly for a certain percentage of trials. We fixed lose-shift at 0.5 based on results of our Bandit task (Figure 2) and empirically fit our lapse parameter, identifying 0.25 as the value that best fit the real Bandit 100:0 data and then explored the consequence of changing the win-stay parameter systematically between 0.5 and 1.0. This modelling revealed that stronger levels of win-stay resulted in higher accuracy of our agent in deterministic environments, but less so in probabilistic environments (Figure 5A). The model simulated behavioural data in 100:0, 90:10, 80:20, 70:30 and 60:40 environments, we only tested the Bandit 100:0 and 80:20 tasks empirically in mice. In the deterministic Bandit 100:0 data, the modelled behaviour closely fit the empirically measured behaviour. Specifically, we observed a strong positive relationship between win-stay and accuracy in the model, and in the win-stay percentage and accuracy across individual mice (Figure 5B).

**Figure 5.**
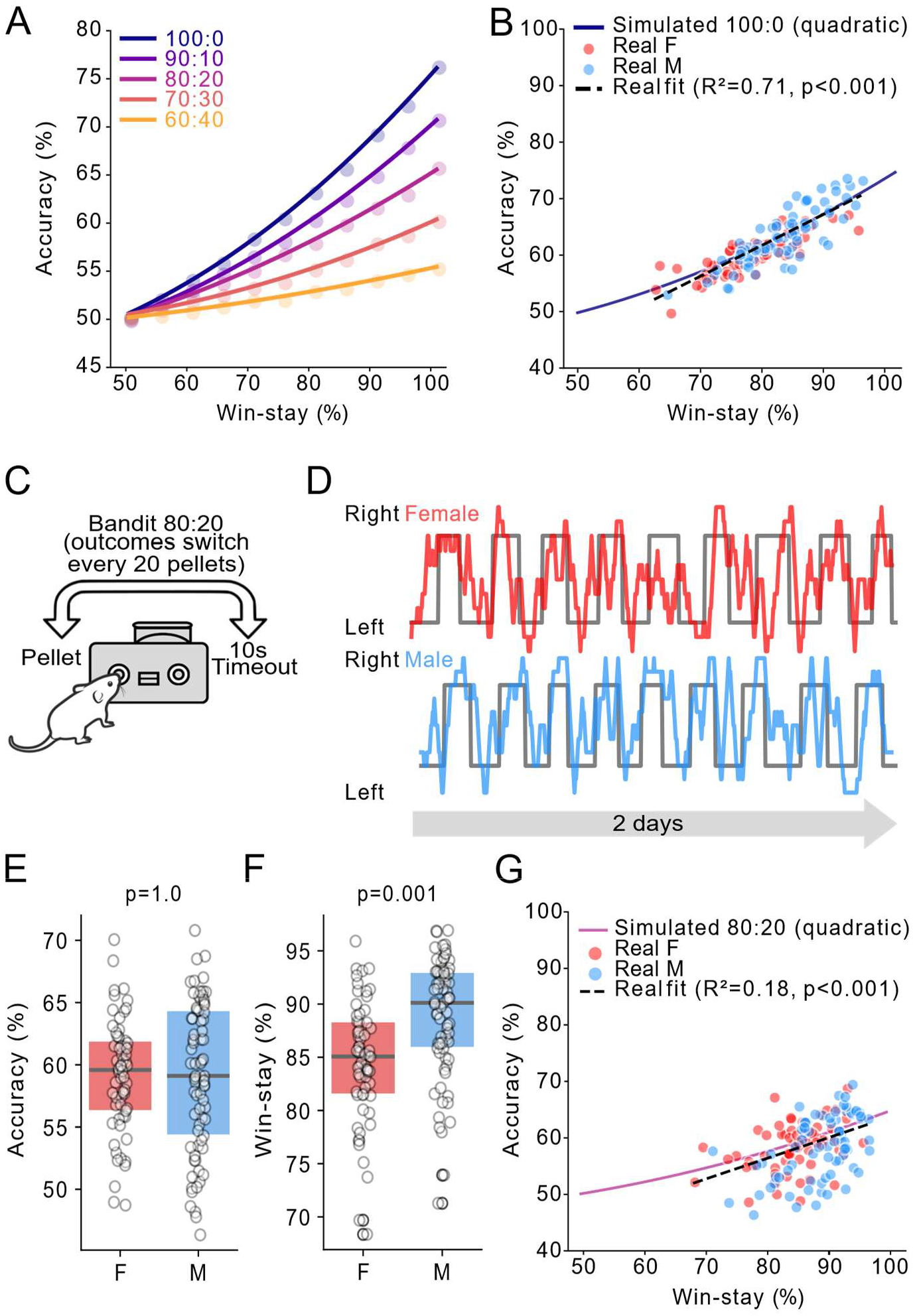
A) Model of win-stay vs. accuracy in the Bandit task, varying poke reward probabilities. B) Fit between Bandit 100:0 behavioural data overlayed with simulated model data. C) Schematic of Bandit 80:20 task. D) Example of mouse behaviour, each datapoint represents the average choice proportion of 10 trials in a rolling window (F red, M blue) with the grey trace indicating the true reward probability in the Bandit 80:20 task. E, F) Accuracy and win-stay probability of females vs males (n = 62F/74M). G) Fit of Bandit 80:20 behavioural data overlayed with simulated model data. P-values were the result of unpaired 2-tailed t-tests, Bonferroni corrected.

Therefore, to empirically test if the stronger win-stay behaviour of male mice failed to boost accuracy in a probabilistic environment, we tested these same mice on a probabilistic variant of the Bandit task for 2 additional days, with probabilities of receiving a pellet or time-out set at 80% vs. 20%, and again switching every 20 pellets (Figure 5C, D). Consistent with our prediction, male mice were no longer more accurate (defined as percentage of pokes on the high probability port) than female mice on this task variant (M = 59.1 ± 0.7%, F = 59.1 ± 0.6%, Figure 5E, unpaired t test *p* = 1.0). We also observed no significant differences in the number of pellets (M = 165 ± 3, F = 174 ± 3 unpaired t test *p* = 0.08) or pokes between male and female mice (M = 593 ± 12, F = 629 ± 10 unpaired t test *p* = 0.08). Male mice still had a higher percentage of win-stay trials than female mice (M = 88.8 ± 0.6%, F = 84.6 ± 0.7%, Figure 5F, unpaired t-test *p* < 0.001), again demonstrating that this appears to be a stable sex difference between male and female mice. We compared this empirical data to the simulated WSLS agent, again observing strong overlap, although the positive relationships between win-stay behaviour and accuracy were not as strong in the modelled or empirical data (Figure 5G).

It is possible that the enhanced win-stay behaviour of male mice reflects a higher motivation or demand for pellets. To test this, we tested these same mice for a final 2 days of a resetting progressive ratio 1 (PR1) task (Matikainen-Ankney et al. 2021). In this task, mice start by earning a pellet with one poke, after which the number of pokes required to earn each pellet increases by one for each earned pellet (Figure 6A). The contingency reset to one poke if they withheld poking for 30 minutes (defined as a “breakpoint”). Mice make multiple “runs” of pellets on this task (Figure 6B).

**Figure 6.**
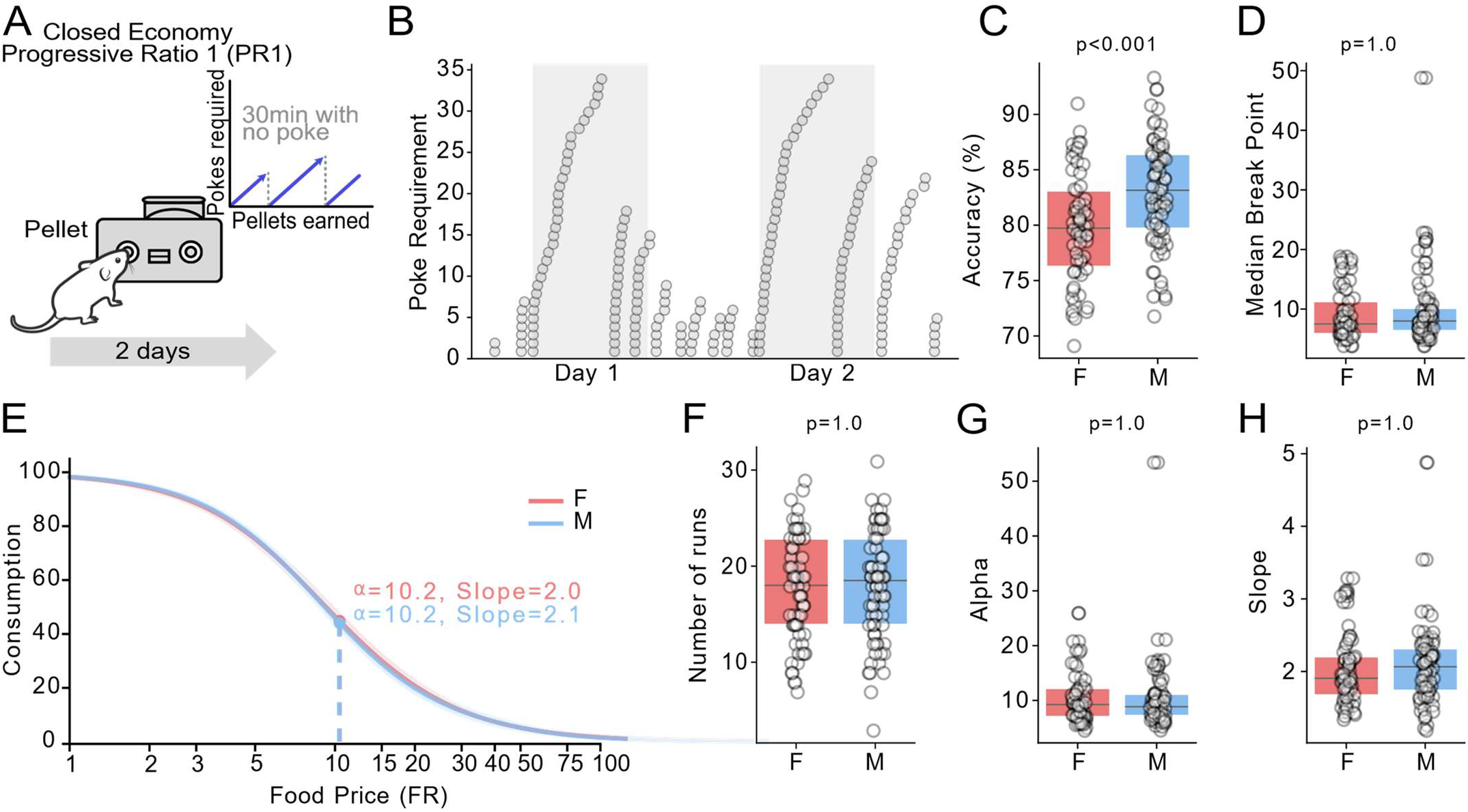
A) Schematic of the progressive ratio 1 task. B) Example of pellets earned over 48 hours, showing “runs” of pellets at increasing poke requirements. C, D) Accuracy and median break point between males and females (n = 62F, 74M). E) Demand curves displaying average consumption of male vs. female mice at each FR (price), with the alpha (price at 50% consumption) and slope of the curve. F) The average number of runs in male and female mice. G, H) Modeled parameters alpha and slope between males and females. P-values were the result of unpaired 2-tailed t-tests, Bonferroni corrected.

There were no significant differences in the number of pellets taken (M = 118 ± 2, F = 117 ± 4, Figure 6C, unpaired t-test *p* = 1.0), or pokes (M = 3663 ± 181, F = 3528 ± 168, unpaired t-test *p* = 1.0). Males had significantly higher “accuracy”, defined as the percentage of pokes on the active port (M = 83 ± 0.6%, F = 79.8 ± 0.6%, Figure 6C, unpaired t-test *p* = 0.001). However, there was no difference in the median breakpoint (M = 9.5 ± 0.7, F = 9.1 ± 0.5, Figure 6D, unpaired t-test *p* = 1.0) or number of “runs” made (M = 18 ± 0.6, F = 17.6 ± 0.7, Figure 6F, unpaired t-test *p* = 1.0). We modelled the PR1 task as a “demand curve”, plotting the number of pellets earned at each ratio (Atalayer and Rowland 2011). The average demand curve of male vs female mice overlapped nearly completely (Figure 6E), and modelled curve parameters of alpha, and beta (slope) were not significantly different (Figure 6G-H, unpaired t-tests, all *p* = 1.0). This argues against our results being explained by gross differences in food demand or willingness to work for pellets between male and female mice.

Our sample size (n = 62F / 74M) is larger than many prior decision-making studies in rodents (Figure 7A). The size of our data set allowed us to model the statistical power of our findings and estimate how likely other researchers would be to replicate our observed sex differences in win-stay behaviour with smaller sample sizes. We simulated experiments of different sample sizes from N = 4 to 60 mice by sampling randomly from our full dataset 1000x at each N (Figure 7B). Each sample was tested for significance, empirically determining the statistical power (the percentage of significant tests) at each N (Figure 7B). The sample size needed to achieve statistical power of 80% was ∼30 mice per group. We also performed an analytical power analysis with the *solve_power()* function in the statsmodels Python package, finding largely overlapping curves (Figure 7C). Importantly, both power curves fell off drastically at lower numbers, with a power of just 20% at 10 mice per group, a common size used in behavioural neuroscience experiments (Figure 7C). We conclude that small sample sizes and under-powered experiments may contribute to a lack of consistent conclusions regarding sex differences in win-stay and lose-shift decision-making strategies between different labs (Button et al. 2013).

**Figure 7.**
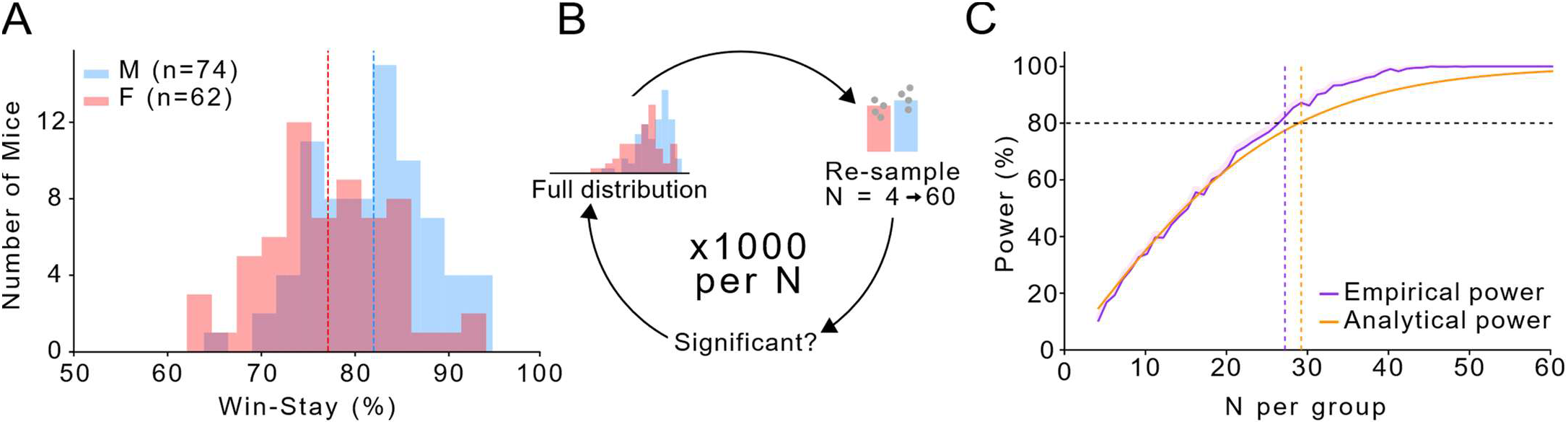
A) The distribution of win-stay % on the Bandit 100:0 task (F red, M blue). Dashed lines represent the mean win-stay % of each sex. B) Schematic of sampling procedure for choosing between 4 and 60 mice per group for empirical power analyses. C) Comparison of empirical vs. analytic power for detecting a sex-difference in win-stay % at different sample sizes.

## Discussion

We aimed to understand sex differences in explore vs. exploit decisions between male and female mice using closed economy, high-throughput, home-cage tasks. Across multiple tasks, males showed stronger exploitative choice patterns, more likely repeating actions that led to prior rewards (ie: win-stay behaviour). Although these effect sizes were small (approximately 5% difference in behaviour), they were highly significant and not explained by differences in food motivation as assessed by pellet intake, breakpoints, and demand curves in a progressive-ratio task. In the deterministic bandit task, but not the probabilistic task, greater exploitation in males was also associated with more efficient foraging, highlighting an interesting interaction between exploitation, efficiency, and environmental certainty. This relationship was supported by behavioural modelling with a win-stay lose-shift (WSLS) agent, suggesting that exploitation produces stronger benefits in more deterministic environments, while exploration may provide additional benefits in less certain environments.

This pattern is consistent with optimal foraging theory, which proposes that animals maximize caloric intake relative to the costs of time and effort (Charnov 1976; Pyke 1984). In patch-based models of foraging, animals must balance exploitation of a specific resource, or “patch” of food, against leaving the patch to explore potentially better options elsewhere. The sex differences in our home-cage tasks align with field observations showing that males spend longer times exploiting food patches than females (Inglis et al. 1996; Garcia et al. 2023), and also with evidence that males persist longer in risky foraging contexts than females (Pellman et al. 2017). Given the similarity between these observations and our home-cage tasks, the relative propensity for exploitation may be a stable sex-difference between male and female mice.

The contrast between deterministic and probabilistic environments clarifies the conditions under which these strategies are adaptive. In the deterministic Bandit 100:0 task, exploitation was advantageous and resulted in greater efficiency of foraging in male mice. However, in the probabilistic Bandit 80:20 task, exploitative behaviour was less beneficial and was no longer associated with significantly greater foraging efficiency in male mice. Although there were only two options in this task, the exploitative strategy that no longer produced higher foraging efficiency, also came at the cost of not exploring other options. This raises the possibility that in more complex environments with multiple options, less exploitative strategies could be more optimal, especially if they enhance exploration and lead to discovery of alternative advantageous options more quickly. Overall, our results support the idea that male and female mice differ in how they solve foraging decisions: males tend to favour exploitation while females rely less on exploitation, consistent with a greater drive for exploration. Neither approach is inherently more optimal across the board, as exploitation produces more optimal foraging in deterministic environments, but not in probabilistic environments (Button et al. 2013). This may have limited their power to detect subtle or more variable sex differences. To address this, we took a high-throughput home-cage approach, which allowed us to acquire a large data set of 136 mice across 4 tasks each. The statistical power afforded by our dataset revealed small but significant sex-differences in foraging efficiency and strategy, which would likely not have been detected in traditionally powered experiments. Specifically, simulations suggested that the significant effects reported here would be difficult to reliably detect with group sizes smaller than 30 mice per sex. It is worth noting that this recommendation is specific for win-stay behaviour. Smaller group sizes may still be sufficiently powered to detect larger behavioural differences, or those that have less biological variability. We make all our raw behavioural data and analysis code freely available online, in part to encourage exploration of power relationships for other metrics in this data.

There are limitations to our study. First, our tasks were trained for a maximum of 3 days in the home cage, which is a relatively short time. While this abbreviated training period had important advantages for throughput, it may have limited our ability to detect other sex-specific differences in strategy that may have emerged with longer training periods. For example in a different set up of the bandit task the International Brain Laboratory tests the duration it takes for mice to become proficient at an 80:20 bandit task, and suggest this takes on average 20 thousand trials (The International Brain Laboratory et al. 2021). Moreover, a study tested one reversal after acquisition on the left poke and finds it takes the mice ∼400 trials to reach an accuracy of 85% after the reversal (Murdaugh et al. 2024). In our tasks mice completed an average of of 859 trials (Bandit 100:0) and 611 trials (Bandit 80:20) where the rewarded side reverses every 20 rewards. Future studies which run these tasks over 6-8 weeks could test when animals may completely learn the task structure and whether their strategy differs with extended experience. For example, mice of both sexes showed a strong influence of win-stay behaviour on their overall strategy in this task, with little evidence that choices further back in time influenced current decisions. Fully trained mice on the bandit tasks could decipher whether males are still less accurate due to an increased win-stay or if more efficient strategies emerge. We also acknowledge that the win-stay lose-shift model we use in this study may not capture all aspects of the mouse behaviour. If mice experienced more training they may make use of additional information about the task structure, and more detailed models may provide more information on the mouse behaviour. Furthermore, this study tested mice using the FED3 device in a single housed environment, this environment could contribute to sex differences that may not translate to other environments or studies using group housing parameters.

Second, although we interpreted the higher win-stay behaviour of males as reflecting greater exploitation of known rewards, this is not necessarily the only explanation. Higher win-stay could reflect differences in learning rate, repetitive behaviours, or sensitivity to rewards between male and female mice. While we explored food motivation and demand with the PR1 task, further experiments may be needed to dissociate these processes more fully. Another explanation could be that female mice are more explorative we however do not directly test exploration in this battery of tests and therefore cannot make this conclusion. A future study disseminating sex differences in exploration would be able to test this hypothesis.

Third, we did not track estrous cycle of the female mice in these experiments, which could potentially contribute to differences in exploitation. Tasks were run across 2 weeks and mice were sampled over random periods, yet the same general trend was seen in multiple tasks over that period, suggesting that estrous cycle was not a primary driver of sex differences. However, formally testing effects of estrous cycle and sex hormones would require dedicated experiments. Fourth, the mice used in this study were non-mutant littermates from 8 lines of mice bred for a study on developmental disorders (WUSMAC project, https://sites.wustl.edu/wusmac/). Although only non-mutant littermates were included in this study, these mice were raised by at least one heterozygous parent with siblings that contained mutations for genes that have been linked to developmental disorders or schizophrenia (parental genotypes given in Table S1). While we cannot rule out an interaction between developmental lineage and sex differences in behaviour, the sex differences on accuracy were directionally consistent across 6 of 8 lines, leading us to conclude that our main result was not specific to their lineage. Finally, all experiments were performed in single-housed C57BL/6J mice in our specific home-cage based tasks. Although this was advantageous for collecting large-scale behavioural data, our results may not necessarily generalize to other strains, housing conditions, or tasks.

We conclude that male mice exhibit slightly but reliably higher levels of exploitation than female mice. These differences were consistent across multiple tasks, were not explained by differences in food motivation, and had distinct behavioural consequences depending on environmental certainty, with greater exploitative bias conferring an advantage in deterministic but not probabilistic settings.

Importantly, both sexes successfully met their energetic needs, indicating that neither strategy was inherently superior, but rather that males and females differ in how they select and deploy decision-making policies. These findings suggest that exploration–exploitation balance represents a behavioural phenotype reflecting underlying computational and neurobiological mechanisms, potentially including differences in reinforcement learning parameters, action repetition biases, or neuromodulatory systems such as dopaminergic signalling. In this context, sex differences in strategy may provide a framework for understanding sex-biased vulnerability in neuropsychiatric disorders characterized by maladaptive exploration or exploitation. More broadly, our results highlight that sex differences in decision-making arise from differences in strategy selection rather than overall performance, emphasizing that multiple behavioural solutions can support successful outcomes in dynamic environments.

## Methods

### Mice and husbandry

C57BL/6J background mice aged 8-16 weeks were individually housed for each experiment on a 12h light/dark cycle. Mice in this study were non-mutant littermates of mice that were bred for the WUSMAC project (https://sites.wustl.edu/wusmac/) and therefore had at least one heterozygous parent and siblings that contained mutations for genes that have been linked to developmental disorders or schizophrenia (parental genotypes given in Table S1). All procedures were approved by the Animal Care and Use Committees at Washington University in St Louis.

### Diets

Grain pellets (20mg grain-based pellets, 10% fat, 25% protein, 64% carbohydrates, 3.35kcal/gm metabolizable energy, Bio-Serv F0163) were provided via the FED3 device on testing days, and standard laboratory chow (5053 - PicoLab® Rodent Diet 20, 13% fat, 24% protein, 62% carbohydrates, 3.02kcal/gm metabolizable energy) was provided *ad libitum* when FED3 was not in the cage. Water was always available *ad libitum*.

### Behavioural testing: FED3

Mice were individually housed and given free access (with no device) to grain pellets and standard chow 3 days before behavioural testing started. No training period occurred prior to the initiation of this experiment. Behavioural testing was conducted using the Feeding Experimentation Device 3 (FED3), a battery-powered, programmable pellet dispenser (Matikainen-Ankney et al. 2021). FED3 devices were placed in the home cage, allowing mice to interact with the device around the clock for multiple days under closed-economy conditions, meaning that all food pellets were earned from the FED3 device. Across all tasks, FED3 logged the timing of nose-pokes, pellet deliveries, and pellet retrievals for offline analysis. Each mouse completed four tasks on FED3 in this study, taking place across two weeks. Bandit 100:0 (3 days) and FR1 (1 day) were completed in the first week, while Bandit 80:20 (2 days) and PR1 (2 days) were completed in the second week. Mice were returned to *ad libitum* chow with the FED3 removed from their cage during the weekend between the two testing weeks.

### Exclusion criteria for behaviour testing

Mice were excluded based on predefined criteria to ensure data quality and animal welfare. Mice were excluded if they did not achieve enough calories in a day defined as 75 pellets across any tasks. Sessions were also excluded in cases of technical issues, including device malfunctions, pellet dispenser jams, or data logging failures. Animals were further excluded if we did not have a full data set of all four behavioural tasks after accounting for session-level exclusions.

### Bandit 100:0

In the 100:0 bandit task, poking on one nose-poke resulted in pellet delivery with 100% probability, whereas poking the other resulted in a 10-second timeout with a white noise stimulus delivered the entire time. These contingencies switched every 20 pellets, such that the previously rewarded port now delivered the 10-second timeout, and the previously unrewarded port delivered a pellet. As in all FED3 tasks, a new pellet could not be earned until the previously delivered pellet was removed from the well. All mice completed this task for 3 days straight, resulting in ∼15-20 contingency switches.

### Fixed-ratio 1 (FR1)

In FR1, the Left nose-poke delivered a pellet, while the other poke had no programmed consequence. After pellet delivery, the mouse was required to retrieve the pellet from the food well before another pellet could be earned, preventing pellet accumulation in the hopper. There was no additional timeout after pellet retrieval, so mice could earn pellets as quickly as they ate them.

### Bandit 80:20

This variant of the Bandit task reduced the probability of reward from 100% to 80% on the high-value port, and increased the probability of reward from 0% to 20% on the low-value port. All trials where a pellet was not delivered resulted in a 10s timeout with white noise stimulus, regardless of whether they occurred on the high or low-value port. Mice completed this task for 2 days straight.

### Progressive-ratio 1 (PR1)

In PR1, mice started by earning a pellet with one poke on the Left side, and for each successive pellet the response requirement increased by one nose-poke. Thus, the second pellet required two pokes, the third required three pokes, etc. Because the task was run continuously in the home cage under closed-economy conditions, a reset rule was used to allow animals to reset the ratio after pauses in responding: if the mouse made no nose-poke on the left for 30 minutes (defined as a “break-point”), the response requirement reset back to 1. In this way, animals completed multiple “runs” across the task, allowing us to obtain repeated measurements of breakpoints and other metrics.

### Behavioural modelling

We modelled behaviour of an agent interacting with FED3 using a win-stay lose-shift (WSLS) strategy. The WSLS agent followed a simple trial-by-trial heuristic: after a rewarded choice, it repeated the same choice on the next trial (“stay”) with a set probability, whereas after an unrewarded choice, it switched with a set “lose-shift” probability. The “lose-shift” probability was fixed at 0.5, while the “win-stay” probability was systematically varied between 0.5 and 1.0. A fixed lapse parameter was also used, such that on 25% of trials the agent dropped this WSLS rule and instead made a random choice. The lapse parameter was obtained by fitting modelled performance to empirical data. The model was run for 100 simulated mice at each probability pair (100-0, 90-10, 80-20, 70-30, and 60-40), with 300 simulated trials per mouse.

### Principal Component Analysis and classification of sex

For dimensionality reduction, we performed principal component analysis (PCA) on behavioural metrics extracted from the FR1 and Bandit100 tasks. The analysis included seven behavioural metrics from the Bandit 100 and FR1 task: Pellets, Pokes, Accuracy, Win-stay, Lose-stay, Retrieval time and Poke time. Retrieval time is defined as the time from the pellet dispensing into the well and the mouse picking it up. Poke time is the time the mouse spends in the nose poke port performing a poke. Before PCA, all behavioural metrics were normalised using z-scores. PCA was then performed, and the first two principal components were used for visualisation. Mice were plotted in PC1–PC2 space and coloured by sex. To identify the behavioural metrics contributing most strongly to the primary axis of variation, PC1 loadings were extracted and ranked by absolute magnitude. The eight variables with the largest absolute PC1 loadings (Retrieval time, Win-stay, Pokes and Accuracy from both Bandit 100 and FR1) were selected for downstream classification analyses.

### Data analysis and statistics

CSV files generated by FED3 were processed with custom python scripts (Python, version 3.6.7, Python Software Foundation, Wilmington, DE). Data wrangling and plotting used Pandas (Mckinney 2010), Matplotlib (Hunter 2007), Pingouin (Vallat 2018), and Seaborn (Waskom 2021) packages. One- or two-way ANOVAs with post-hoc t-tests or paired or unpaired t-tests were used to compare groups as appropriate, using Bonferroni corrections to control family-wise error rate. P-values < 0.05 were considered significant. Data sets are presented as mean ± SEM. The number of animals per experiment is listed as n=number of animals. ChatGPT and Google Gemini were used for assistance with coding, but the Authors take full responsibility for all analysis code. All data and analysis and visualization code are available at https://github.com/KravitzLab/Murrell2026.

### Demand curve modelling for the PR1 task

PR1 data were modelled as a behavioural economic demand curve by quantifying the number of pellets earned as a function of fixed-ratio requirement (i.e., price) (Atalayer and Rowland 2011). Demand curves were fit separately for each mouse and normalized to maximal consumption at Q0, the theoretical consumption at zero cost. The parameter α, representing the food price at which consumption declined to 50% of its maximal value, provides an index of demand elasticity. We also report the slope at α, which reflects how sharply consumption falls around this transition point and thus provides an index of cost sensitivity across the dynamic range of the curve.

### Power analyses

We performed both empirical and analytical power analyses on our data. For the empirical testing, we randomly sampled win-stay probabilities in groups of between 4 and 60 males and females 1,000 times. Each sample was tested for significance, noting the percentage of the 1000 samples that achieved significance. This was plotted as the empirically derived statistical power. We also used the *solve_power()* method from statsmodels to analytically determine the power at the same sample sizes between 4 and 60 subjects per group.

## Acknowledgements

This work was supported by the NIH R01DK136810 (AVK), DK138131 (AVK), RM1MH138313 (JDD, SEM, AVK), DA049924 (MCC), NIH P50 HD103525 (JDD) and SFARI 734069 (JDD),and the Taylor Family Institute for Innovative Psychiatric Research, Washington University School of Medicine. We thank the Animal Behaviour Subunit of the IDDRC at WashU for access to procedural space, and the Department of Neuroscience NeuroTech Hub for technical assistance.

## Code and data availability

All data and code to generate figures are available at: https://github.com/KravitzLab/Murrell2026.

## Notes

https://github.com/KravitzLab/Murrell2026

### Competing Interest Statement

The authors have declared no competing interest.

### Summary of Updates

Updated two typographical errors in the figures.

